# Perception of and anxiety about COVID-19 infection and risk behaviors for spreading infection: An international comparison

**DOI:** 10.1101/2020.07.30.228643

**Authors:** Akihiro Shiina, Tomihisa Niitsu, Osamu Kobori, Keita Idemoto, Tasuku Hashimoto, Tsuyoshi Sasaki, Yoshito Igarashi, Eiji Shimizu, Michiko Nakazato, Kenji Hashimoto, Masaomi Iyo

## Abstract

To control the spread of the newly developed corona viral infection diseases (COVID-19), people’s appropriate precautionary behaviors should be promoted. We conducted a series of online questionnaire survey, to gather a total of 8,000 citizen’s responses on March 27–28, 2020 in Japan and April 17–21 in the UK and Spain. Compared to Japan, the knowledge and anxiety level and the frequency of precautionary behaviors were higher in the UK and Spain. Participants with infected acquaintances were more concerned about COVID-19. However, participants in the UK rarely wore a medical mask. Participants in the UK and Spain were eager to get information about COVID-19 compared to those in Japan. The participants in Spain tended not to trust official information and to believe specialists' comments instead. The urgency of the spread of COVID-19, cultural backgrounds, and recent political situations appear to contribute to the differences among countries revealed herein.

## Introduction

In December 2019, a novel species of coronavirus which causes very specific and critical pneumonia was identified: SARS-CoV-2. This virus expanded all over the world within several months, leading to millions of infections. Considering this deteriorating situation, the World Health Organization (WHO) declared a pandemic state on March 11, 2020. Citizens in many countries are now facing the risk of the very serious disease caused by SARS-CoV-2, i.e., COVID-19.

Facing uncertain situations can increase people's anxiety levels, especially when there is potential risk for mortality. This anxiety may lead both healthy and vulnerable individuals to engage in behaviors designed to protect them from contracting the virus [1]. Cao et al. reported that roughly one-fourth of the college students evaluated in China described feeling at least mild anxiety because of the COVID-19 outbreak [2]. The fear about COVID-19 is expected to have a major impact on public mental health [3]. Jones and Salathé reported that the engagement in protective behaviors varies from person to person, and may be affected by several factors [4].

According to the WHO [5], SARS-CoV-2 is transmitted during close contact through respiratory droplets (such as coughing) and by fomites. For the prevention of the transmission of SARS-CoV-2, the WHO [5] continues to recommend performing frequent hand hygiene, using respiratory protection, regularly cleaning and disinfecting surfaces, maintaining physical distances, and avoiding people with fever or respiratory symptoms. Based on the WHO's recommendations, most countries have taken steps to address the local circumstances of the epidemic as well as the accompanying economic downturn. However, the development of further strategies to achieve greater control of the current pandemic situation is required.

We have been concerned about the differences among countries regarding the populations' cognitional and behavioral patterns as well as attitudes toward information sources in relation to the anxiety about the COVID-19 pandemic. Knowing these differences will contribute to our understanding of the patterns of epidemic-related anxiety and behaviors, and it will help optimize future policies for preventing the second wave of the epidemic.

## Materials and Methods

We conducted a series of web-based cross-sectional surveys to determine the international differences in their populations' perceptions and behaviors in the current COVID-19 risk situation. We asked Cross Marketing Inc. to recruit a total of 8,000 individuals for this series of surveys, which were conducted with two stages. In the first stage, we conducted the survey of 4,000 residents of Japan. The results were independently analyzed and published [6]. These data are dealt with as a reference in the present report. In the second stage, we conducted the survey of 2,000 individuals in the UK and 2,000 individuals in Spain. In this report, we present the data from the UK and Spain combined with those from Japan.

Participants had to be over 20-year-old. People who had been diagnosed with COVID-19 infection were excluded from the study.

We adopted the questions used in a previous report [4] but included new questions about anxiety levels regarding symptomatic aggravation and virus transmission. In addition to the demographic information, the questionnaire included several items covering the respondent's level of fear and anxiety about COVID-19-related issues, and the frequency of the respondent's media exposure, trust in each media source, and frequency of anti-infection behaviors. The participants provided answers on scales from 1 (none/never) to 9 (extremely/strongest) for the items regarding their understanding of the symptoms, preventive methods, health management and consulting services when infected, levels of fear and anxiety, and the frequency of anti-infection behaviors. Items regarding the frequency of media exposure and degree of trust in each media source were rated on scales from 1 (almost none/not at all) to 5 (very/greatly). We also asked the UK and Spain participants about the frequency of hand washing in their usual life. The questionnaire is presented as a Supplementary Table. We did not ask the Japanese participants about the frequency of handwashing because a past survey had done so [7].

We analyzed the gathered data using SPSS for Windows, ver. 24 (IBM, Armonk, NY). We used the χ^2^-test for data with nominal scales, an analysis of variance (ANOVA) with the Games-Howell test for parametric data, and the Kruskal-Wallis test with Dunn-Bonferroni's correction for non-parametric data. The level of significance was set at p<0.05.

Participants were informed by electric letter that their participation was voluntary. We did not gather any personal information about the responders. All respondents were taken to agree to participate if they sent in their answer form. The participants were rewarded according to the regulations of Cross Marketing Inc. The whole study protocol was approved by the Ethics Committee of Chiba University Graduate School of Medicine and the Ethics Committee of the International University of Health and Welfare before implementation.

This study was conducted with a management grant provided by Japan's Ministry of Education, Culture, Sport, Science and Technology [MECSST] to Chiba University Graduate School of Medicine and by a grant from the Japan Society for the Promotion of Science [JSPS] KAKENHI [to T.N., no. JP19K08066]. The MECSST and JSPS had no role in or control of the execution of this study.

## Results

Between March 27 and 28, 2020, a total of 4,000 participants in Japan stratified by age (20s, 30s, 40s, 50s, and ≥60 years) and gender (400 in each group) took part in this study. Between April 17 and 21, 2020 a total of 2,000 participants in the UK and 2,000 participants in Spain stratified by age (20s, 30s, 40s, 50s, and ≥60 years) and gender (200 in each group) took part in the study. The concurrent situations in each country are presented in Table 1.

**Table 1.** The participants’ demographic characteristics

|  |  | Japan |  | UK |  | Spain |  |
| --- | --- | --- | --- | --- | --- | --- | --- |
|  |  | n | % | n | % | n | % |
| Age | 20s | 791 | 19.9 | 393 | 19.8 | 396 | 19.8 |
|  | 30s | 793 | 19.9 | 398 | 20.0 | 400 | 20.1 |
|  | 40s | 800 | 20.1 | 399 | 20.1 | 399 | 20.0 |
|  | 50s | 799 | 20.1 | 399 | 20.1 | 400 | 20.1 |
|  | ≥60s | 798 | 20.0 | 398 | 20.0 | 400 | 20.1 |
|  | Total | 3981 | 100 | 1987 | 100 | 1995 | 100 |
| Gender | Male | 1984 | 49.8 | 993 | 50.0 | 997 | 50.0 |
|  | Female | 1997 | 50.2 | 994 | 50.0 | 998 | 50.0 |
|  | Total | 3981 | 100 | 1987 | 100 | 1995 | 100 |
| Educational background | Junior high school/Secondary school | 106 | 2.7 | 258 | 13.0 | 379 | 19.0 |
|  | High school/A-level or equivalent | 1223 | 30.7 | 601 | 30.2 | 242 | 12.1 |
|  | Diploma course or vocational school | 876 | 22.0 | 306 | 15.4 | 526 | 26.4 |
|  | University degree or above | 1776 | 44.6 | 822 | 41.4 | 848 | 42.5 |
|  | Total | 3981 | 100 | 1987 | 100 | 1995 | 100 |
| Infected family or relatives | Yes | 17 | 0.4 | 498 | 25.1 | 779 | 39.0 |
|  | No | 5909 | 95.1 | 1183 | 59.5 | 939 | 47.1 |
|  | Uncertain | 177 | 4.4 | 306 | 15.4 | 277 | 13.9 |
|  | Total | 3981 | 100 | 1987 | 100 | 1995 | 100 |
| Infected persons in your workplace | Yes | 45 | 1.1 | 342 | 17.2 | 377 | 18.9 |
|  | No | 3744 | 94.0 | 1263 | 63.6 | 1278 | 64.1 |
|  | Uncertain | 192 | 4.8 | 382 | 19.2 | 340 | 17.0 |
|  | Total | 3981 | 100 | 1987 | 100 | 1995 | 100 |

We first excluded invalid answers from the analysis because there might be some participants who gave non-serious answers considering that participating in the survey would be rewarded. Participants whose answer to all 30 of the items in section C, D, E, and F excepting number of hand washing was '1' were excluded. As a result, 19 participants in Japan, 13 in the UK, and 5 in the Spain were excluded from the analysis. The remaining 7,963 participants were subjected to the analyses.

The participants' demographic data are presented in Table 2. Regarding educational background, the participants in Japan had a higher percentage of university degrees than the participants in the UK and Spain (χ^2^-test, Pearson χ^2^ = 674.390, df = 6, p<0.001, adjusted residual for university degree or above of the participants in Japan = 2.4). Compared to the participants in the UK and Spain, those in Japan had a lower percentage of having infected people among their family or relatives (χ^2^-test, Pearson χ^2^ = 2055.999, df = 4, p<0.001, adjusted residual for participants in Japan for whom this answer was 'yes' = −38.3), or at their workplace (χ^2^-test, Pearson χ^2^ = 1139.305, df = 4, p<0.001, adjusted residual for participants the answer 'yes' in Japan = −18.6).

**Table 2.**
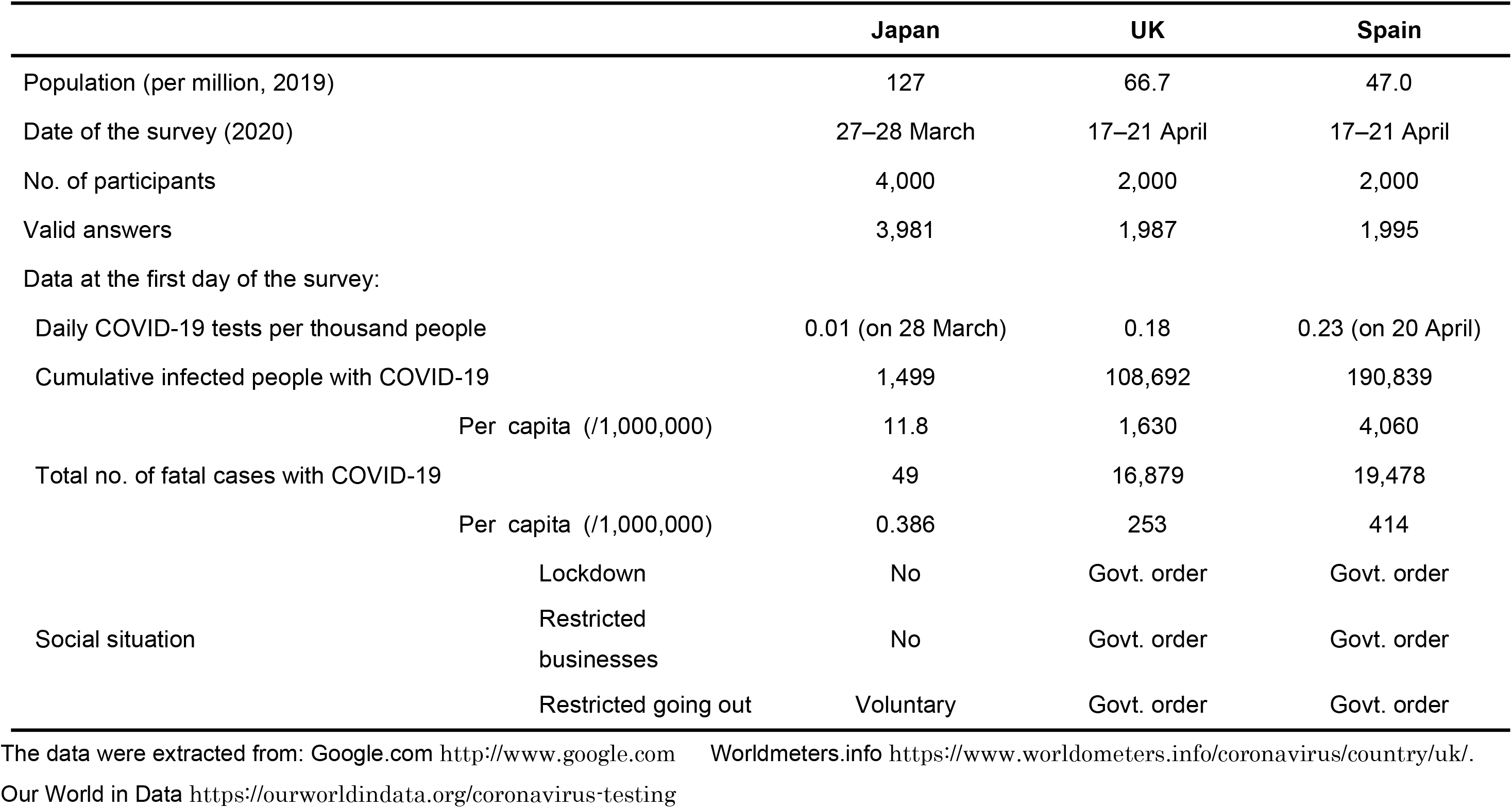
Situations in each nation at the time of survey

Regarding the level of knowledge about COVID-19, the results of this series of surveys revealed definite and significant differences among the three countries, as illustrated in Figure 1. The participants in the UK and Spain had a deeper understanding of COVID-19 than the participants in Japan (ANOVA with Games-Howell test).

**Fig. 1.**
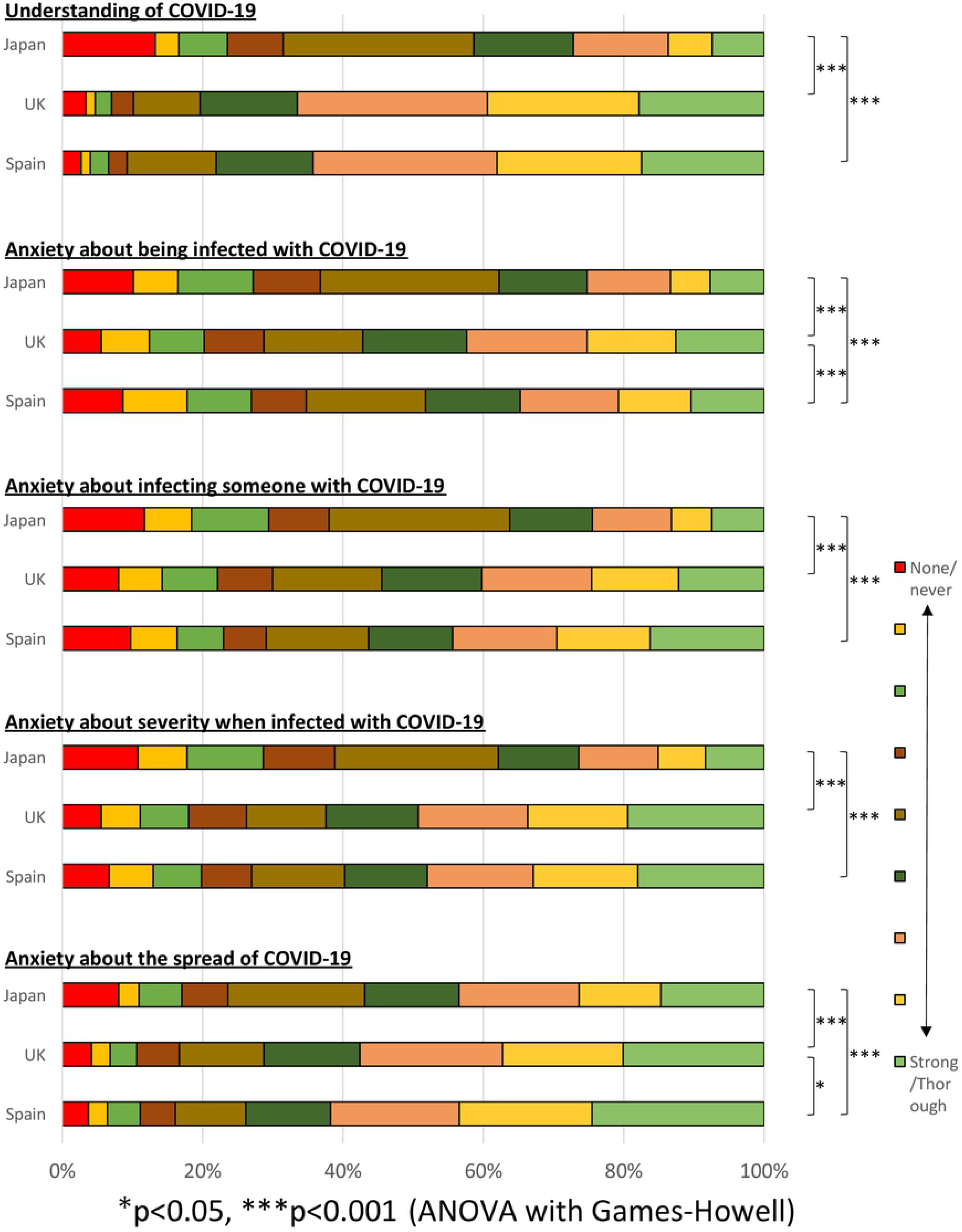
Knowledge and anxiety about COVID-19 in each country The level of each item was graded from 1 to 9, with larger values indicating higher levels of that factor.

Regarding anxiety about COVID-19, there were also significant differences among the countries (Fig. 1). The participants in Spain were more anxious about being infected with COVID-19 than those in Japan, but less anxious than those in the UK. The participants in Japan were less afraid of infecting others with COVID-19 than those in the UK and Spain. Regarding their anxiety about the severity of the disease once infected, the participants in Japan were less concerned than those in the UK and Spain. Concerning the spread of COVID-19, the participants in Spain were the most anxious, followed by those in the UK and then Japan.

We examined the frequency of access to and the credibility of a variety of sources of information about the virus and pandemic, and the participants' survey responses revealed some country-specific characteristics (Fig. 2). The participants in Japan reported having significantly less frequent access to any form of information compared to those in the UK and Spain. Remarkably, the proportion of participants in Japan who said that they had never had any access to official announcements, radio, or specialists was twice as high as that of the other two countries. The participants in Spain had more frequent access than those in the UK to information sources such as the government, a social network, radio, friends and neighbors, and specialists.

**Fig. 2.**
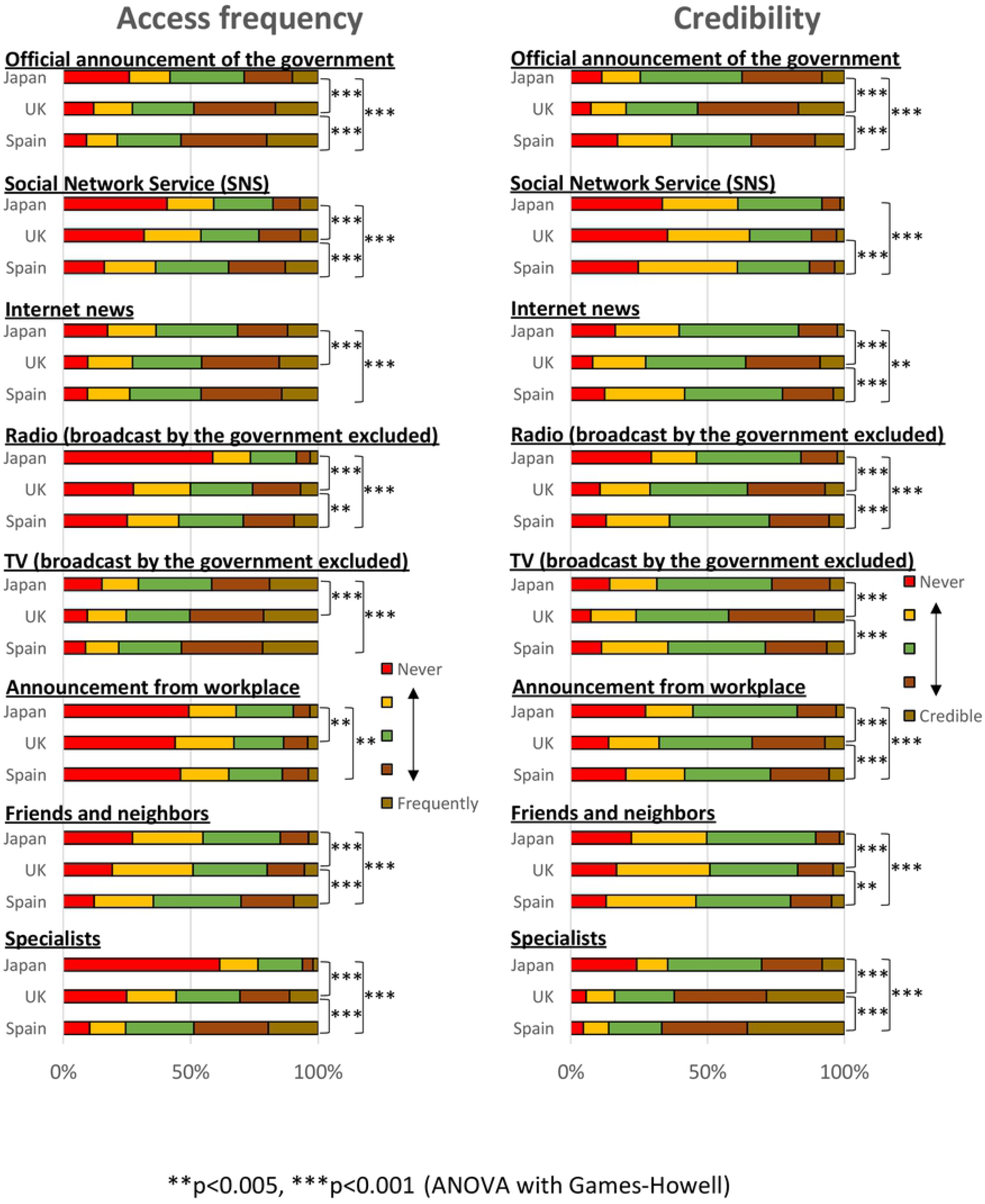
Attitude toward information sources in each country The level of each item was graded from 1 to 5, with larger values indicating higher frequency/credibility.

Regarding the credibility of the COVID-19 information sources, the participants in Japan were unlikely to trust most of the types of information sources compared to those in the UK and Spain. The participants in Spain had less trust in the information from the government compared to the participants in Japan and the UK. Social networks and friends and neighbors were not deemed a credible information source in any of the three countries, but the participants in Spain reported slightly more trust in these sources compared to those in Japan and the UK.

We compared the number of daily handwashing among countries. The range of handwashing was 4–5×/day in all three countries (data not shown). A Kruskal-Wallis test with Dunn-Bonferroni's correction revealed that the participants in Spain washed their hands significantly fewer times over the course of a day than the participants in the UK (p<0.001), but more times than those in Japan reported in the past survey (p<0.001).

The questionnaire answers regarding precautionary behaviors are illustrated in Figure 3. Among the active behaviors, the patterns of handwashing and using disinfectant showed significant differences among the countries, but the elements of these differences were complex. For example, some UK respondents washed their hands very frequently, although there were also more respondents in the UK who never washed their hands compared to Spain and Japan. We also observed that 57.7% of the participants in the UK never wore a medical mask whereas 16.1% and 9.5% of those in Spain and Japan did, respectively. The participants in Japan were less likely to engage in avoidance behaviors compared to those in the UK and Spain. Notably, the participants in Japan were far less likely to avoid school or work and less likely to avoid using public transport than the other countries' respondents. The majority of the participants in the UK and Spain reported that they rarely or never went to school or work, or used public transport.

**Fig. 3.**
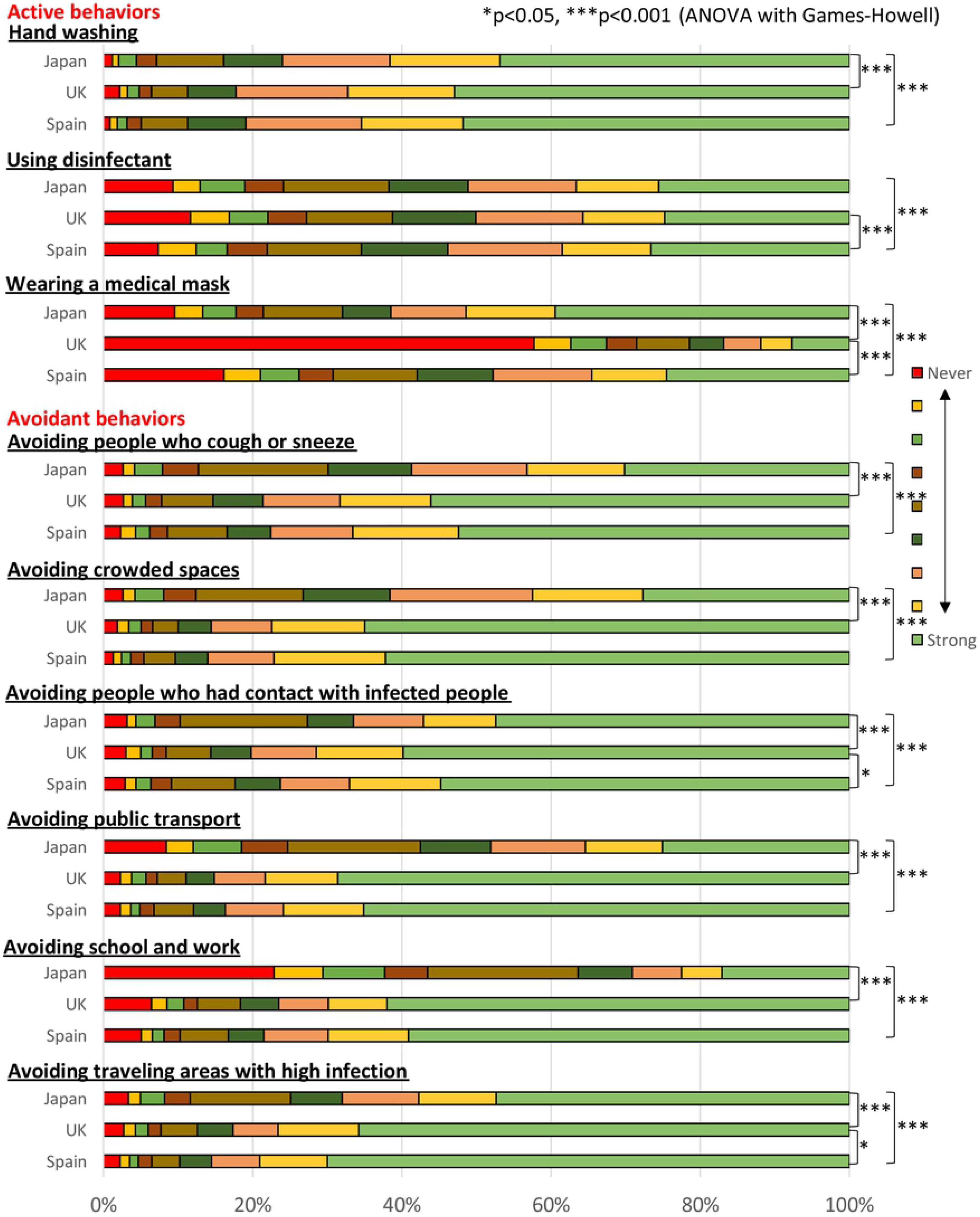
Frequency of precautionary behaviors in each country The level of each item was graded from 1 to 9, with larger values indicating higher levels of that factor.

As an additional analysis, we compared the data of the degree of knowledge, anxiety, frequency and credibility of virus information, and precautionary behaviors between the participants who had acquaintances who were affected by the virus and those who did not. Almost every item in the questionnaire showed a significant difference between these two groups.

We reanalyzed the differences in the items presented above between countries by examining only the participants who did not have an infected acquaintance. The influence of eliminating the answers of the participants with an infected acquaintance was limited.

## Discussion

We conducted an international online series of questionnaires involving a total of 8,000 individuals to investigate their knowledge, anxiety, protective behaviors, and access to information regarding COVID-19 during early Spring 2020. At the time of this investigation in Japan, the spread of COVID-19 was limited to several areas, but a few days after the questionnaire was administered, the number of infected people rocketed upwards and the government declared a state of emergency (but in specific prefectures, it was announce on 7th April [8] that resident not leave their homes without an urgent need to do so). After the end of May, the government began to gradually loosen the restrictions [9].

On the other hand, during the period of the questionnaire's administration in the UK, the spread of the infection there was much more severe. Residents were required to stay home unless there was an essential need to go out. Outgoing business activities were prohibited with few exceptions, until the government loosened the restrictions on 11th May, 2020 [10]. The number of deaths due to COVID-19 increased linearly during the questionnaire period.

In Spain, a lockdown had been in place since 14th March, 2020 with very strict rules. Citizens were forbidden from leaving their homes unless they had to in order to get food or medical care, or walk their dog. Very recently, there is a plan to loosen this regulation in light of the dropping number of cases of new infection [11]. Spain's government requires its population to wear a face mask on public transport despite the relaxed lockdown [12].

The present questionnaire revealed that the participants in the UK and Spain had a deeper understanding of COVID-19 than those in Japan. The participants in the UK and Spain also showed greater anxiety about each item compared to those in Japan. The frequency of access to sources of COVID-19 information was also significantly higher in Spain and UK than Japan. The differences in the attitude of the three countries' residents toward COVID-19 can be explained mainly by the differences in the current pandemic situation in each country. A greater spread of infection has increased both the levels of anxiety and the knowledge of the virus, and it drives the need for more precise information.

Similar international surveys have been conducted recently by other organizations. Dryhurst et al. [13] conducted an international comparison study of the perception of the risk of COVID-19, and they reported that the respondents in the UK had a higher level of risk perception than those in Japan and Spain. These results are consistent with our present findings. Gallup International also conducted a series of international surveys, and according to the results, Japanese were less likely to be anxious about catching COVID-19 than individuals in the UK, whereas more Japanese were very or somewhat scared about COVID-19 at the same time as our questionnaire period [14]. Our results are generally consistent with those of the other concurrent surveys.

Concerning the respondents' precautionary behaviors, we observed differences in the response patterns of active behaviors. Our findings indicated that the respondents in the UK and Spain more frequently their wash hands compared to the respondents in Japan before the pandemic. To the best of our knowledge, there are no other reports presenting an international comparison of three countries regarding the frequency of handwashing. The reason why the respondents in the UK wash their hands significantly more frequently than those in Spain remains to be clarified.

We found notable differences in mask-wearing among the countries. The UK participants were far less likely to wear a mask. The difference cannot be explained by the degree of infection spread; rather, it may be because of the government policy in each country as well as the cultural background [15]. The WHO [15] did not recommend that healthy people wear a face mask unless there are rational conditions. The UK government has not encouraged its citizens to wear a face mask, since there is scarce evidence of the effectiveness of the mask for preventing infection [15][16]. In contrast, it is well known that Japanese people are willing to wear a mask in part because there are many individuals suffering from spring allergies. Approximately 40% of Japanese university students wear a mask in the spring, according to a previous survey [17]. The high percentage of mask-wearing in Japan before a rapid increase in symptomatic infected cases might have contributed to a reduction in the number of infectious cases, because wearing a mask potentially can prevent splash infection by blocking saliva containing the virus, while not blocking the coronavirus itself [18]. However, a recent report indicated that many transferences of SARS-CoV-2 occur from presymptomatic carriers [19]. Another recent study described the efficacy of medical mask-wearing by asymptomatic individuals as well as lockdown at the population level [20].

We observed that the participants in Japan were significantly less likely to engage in any of the avoidance behaviors examined compared to those in Spain and the UK. These results were are similar to those of the other surveys conducted concurrently [14]. Going outside one's residence and working outside were seriously restricted in the UK and Spain at the time of our questionnaire, while in Japan, people only in specific regions were advised (not mandated) to stay home. Workplaces were not closed in many parts in Japan at the time of this study. These situational discrepancies contributed to the differences in social activities among the countries.

A survey of Italian subjects indicated that a higher level of knowledge was positively associated with the acceptance of strict mitigation measures such as lockdown [21]. If this finding is applicable to people in other countries, people in the UK and Spain would be more supportive of national lockdown policies compared to Japanese because the former populations have higher risk perception and anxiety according to our present findings. The UK respondents reported a high level of trust in official information, and this is consistent with the report by Gallup International that the UK respondents were likely to believe that their government was handling the coronavirus issues well [14]. However, according to our results it seems that the study population in Spain tended not to trust official information.

The Spanish respondents to our questionnaires were least likely to trust the government, followed by Japan; instead, they were likely to trust specialists as a source of information, while the Japanese hardly trusted any source of information about the pandemic. Differences in the basic trust of officials because of cultural factors as well as recent political conflicts may have contributed to these results. It was reported very recently that an alarming number of individuals in England believe conspiracy theories about COVID-19, and these theories are associated with less adherence to official guidelines for precautionary behaviors [22]. On the contrary, we observed that the questionnaire respondents who had an acquaintance who was infected with the virus were likely to trust official information about COVID-19. Considering these results, the residents of Japan are potentially vulnerable to conspiracy theories in part because of the rare experience of an acquaintance's infection, although there is no evidence suggesting the spread of COVID-19 conspiracy theories in Japan.

There may be several limitations in this study. It was conducted from March to April in 2020. The time point of the survey differed between Japan and the other countries, and this may have influenced the results. In addition, since a series of web-based questionnaires was used, we cannot eliminate the possibility of selection bias among participants. We gathered the responses of only people who were willing to complete online questionnaires.

